# Functional precision oncology for follicular lymphoma with patient-derived xenograft in avian embryos

**DOI:** 10.1101/2023.06.22.546045

**Authors:** Manon Zala, Boris Lipinski, Clélia Costechareyre, Loraine Jarrosson, Romain Teinturier, Edith Julia, Marjorie Lacourrège, Aurélie Verney, Jérôme Guitton, Alexandra Traverse-Glehen, Emmanuel Bachy, Gilles Salles, Sarah Huet, Laurent Genestier, Valérie Castellani, Céline Delloye-Bourgeois, Pierre Sujobert

## Abstract

**Background:** Follicular lymphoma (FL) is an incurable B-cell malignancy that constitutes a quarter of all lymphomas. Although RCHOP immuno-chemotherapy induces high rates of complete remission, almost all FL patients experience multiple relapses post-treatment. The limited understanding of treatment response heterogeneity is due to the absence of *in vitro* or *in vivo* experimental models, primarily because tumor cells heavily rely on their microenvironment to survive. In this study, we present an innovative xenograft model of primary FL cells in avian embryos, circumventing these limitations.

**Methods:** We developed the FL-AVI-PDX model by transplanting 20 biopsy FL samples, including good (n=11) and poor clinical responders (POD24, n=9), into chicken embryos. Each set of embryos was treated with RCHOP or vehicle intravenously. We evaluated the effect of immuno-chemotherapy on tumor volume by light sheet microscopy and on tumor biology by transcriptomic analysis at the single-cell level.

**Results:** We successfully engrafted all samples in avian embryos. We found that RCHOP treatment *in ovo* led to tumor volume reduction, which predicted progression-free survival in multivariate analysis, demonstrating the model’s capacity to capture clinical heterogeneity at the patient level. The FL-AVI-PDX model also provided a unique opportunity to analyze the transcriptomic impact of RCHOP on FL cells using single-cell RNA sequencing. We identified a signature of 21 genes upregulated after RCHOP exposure, displaying significant intra-tumoral heterogeneity. As a proof of concept, we validated the functional involvement of *BAX*, a gene from the RCHOP-induced signature, as a critical effector of immuno- chemotherapy *in vitro* and in avian embryos.

**Conclusions:** The FL-AVI-PDX model is a platform for functional precision oncology in primary FL cells that captures both inter- and intra-patient heterogeneity of clinical response to a complex therapeutic regimen. It offers a unique opportunity to better understand FL biology, opening perspectives for the development of new drugs.

## Introduction

Follicular lymphoma (FL) is the second most frequent type of lymphoma. It typically presents as an indolent disease, allowing for a “watch and wait” approach to be taken with some patients who are minimally symptomatic (Ardeshna *et al*., 2003). However, most patients will eventually require treatment. The standard of care is a combination of an anti-CD20 monoclonal antibody (rituximab or obinituzumab) with chemotherapy (such as cyclophosphamide, hydroxyadriamycine, vincristine and prednisone (CHOP) or bendamustine) followed by a 2-year maintenance with anti-CD20 (Salles, 2020). Despite impressive results achieved with these treatments (median progression free survival of 10.5 years, 10-year overall survival of 80%) (Bachy *et al*., 2019), relapse invariably occurs, and lymphoma remains the primary cause of death for these patients (Sarkozy *et al*., 2019). It is believed that relapses are caused by cells that are resistant to chemotherapy, but there is currently no data that documents intratumoral heterogeneity regarding treatment response.

Of note, the outcome of FL patients is highly heterogeneous, with a particularly poor prognosis for 20% of the patients experiencing disease progression during the first 24 months of initial immunochemotherapy (POD24) (Casulo *et al*., 2015). Despite numerous efforts to prospectively identify these patients through analyzing driver mutations (Pastore *et al*., 2015) or gene expression (Huet *et al*., 2018), none have been successful in personalizing first-line treatment. An alternative approach, known as functional precision oncology, which measures the response of living cells to perturbations, has the potential to improve the prediction of response to therapy (Letai, Bhola and Welm, 2022). However, this type of analysis is limited in FL due to the lack of an adequate model, as primary FL cells cannot be grown *in vitro.* A few patient-derived xenograft (PDX) have been established in immunocompromised mice, but these models are time consuming and not suitable for drug testing in mutliple replicates (Townsend *et al*., 2016). Hence, development of new FL models remains an unmet need to improve patients outcome.

Avian embryos have been recently shown to provide a supportive environment for the growth of human cancer cells, including neuroblastoma, triple negative breast cancer, and melanoma (Delloye-Bourgeois *et al*., 2017; Jarrosson *et al*., 2021, 2023). Furthermore, as only a small number of cells are needed for each engraftment, multiple experimental conditions can be tested without any pre-amplification of the patient sample thus allowing statistically powerful analysis. In addition, drugs can be administered intravenously in the embryos, making it possible to test drug sensitivity by measuring tumor volume using light sheet microscopy or analyzing exposed tumor cells through single-cell RNA sequencing (scRNAseq). Here we provide evidence that a new model of FL using PDX in avian embryos (FL-AVI-PDX) captures interpatient heterogeneity regarding treatment response. Additionally, scRNAseq revealed a robust signature of RCHOP induced genes, and an unexpected heterogeneity among lymphoma cells exposed to RCHOP. To validate functionally these findings, we demonstrate that the use of venetoclax to target *BAX*, one of the RCHOP induced genes, could enhance the effects of RCHOP.

## Results

### Establishment of the patient-derived xenograft avian model of follicular lymphoma (FL- AVI-PDX)

FL cells are highly dependent on their microenvironment to survive (Scott and Gascoyne, 2014), which may explain why previous attempts to grow these cells *in vitro* have failed. We hypothesized that signals instructing the hemogenic endothelium to generate hematopoietic stem cells in the avian embryos could offer a supportive microenvironment. To test this hypothesis, we injected FL cells in the aorta-gonado-mesonephros (AGM) embryonic region where hematopoietic stem cells emerge (Jaffredo *et al*., 1998). To that end, mechanically dissociated and viably cryopreserved FL samples from diagnostic biopsies were labelled with carboxyfluorescin succinimydil ester (CFSE) in order to be followed *in vivo* and subsequently implanted within the emerging AGM region of series of E2 (Hamburger and Hamilton stage HH15) avian embryos (Figure 1A, Supp. Table 1). All of the primary FL samples tested (n=20) successfully grew in the AGM with 98% (86-100%) of avian embryos with detectable CFSE+ cells 24h after graft (Supp. Table 2). These findings suggest that the emerging AGM region of avian embryos supports primary FL cells survival.

**Figure 1:**
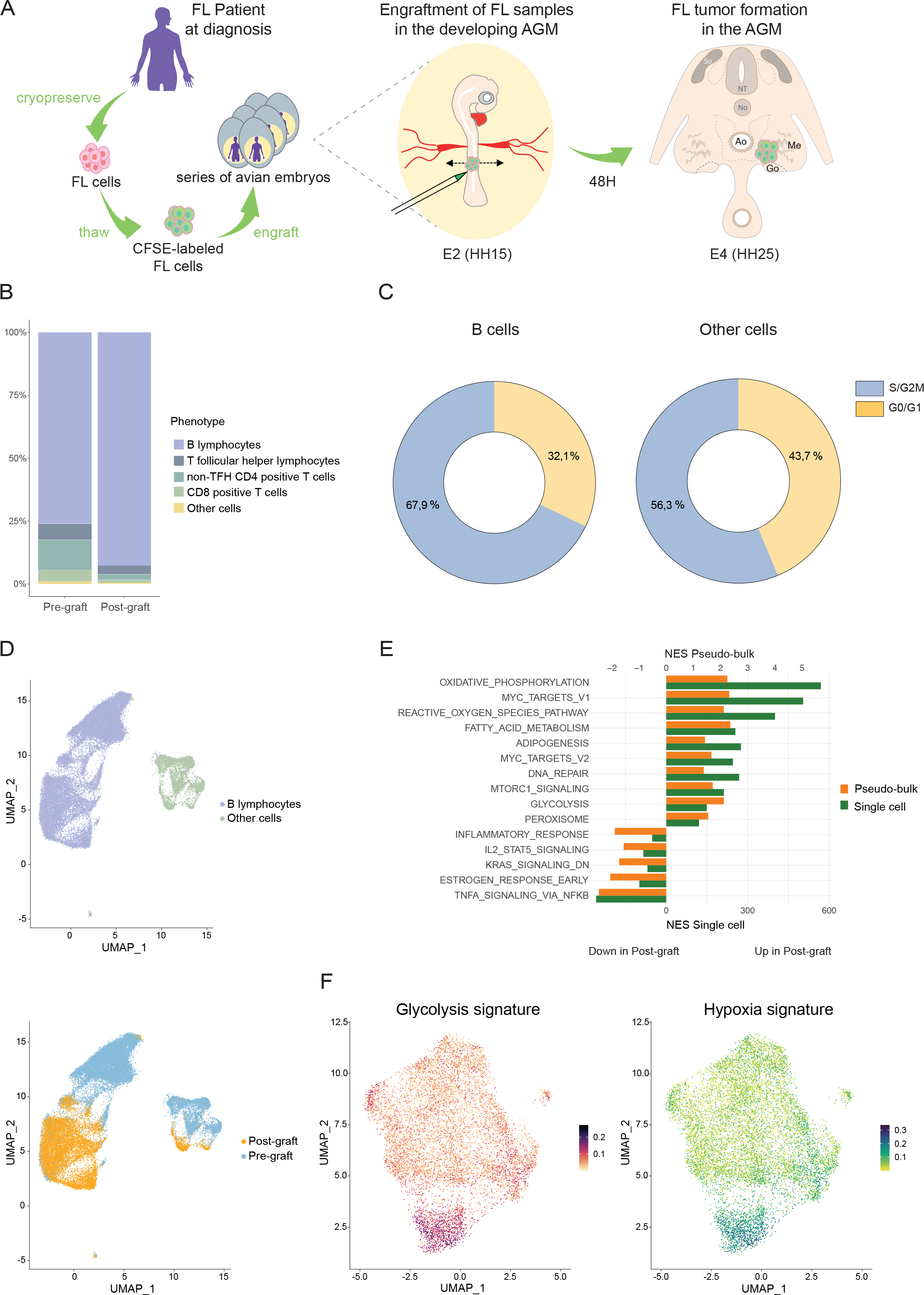
Establishment of the patient-derived xenograft avian model of follicular lymphoma (FL-AVI-PDX). **(A)** Schematic representation of the FL-AVI-PDX model. **(B)** Analysis of cell composition before (pre-graft) and after (post-graft) graft based on scRNAseq data. **(C)** Proportion of S/G2/M or G0/G1 cells among B cell or non-B cells in pre-graft samples. **(D)** Uniform Manifold approximation and projection (UMAP) representation of the unsupervised analysis of scRNAseq data from fourteen patients. Individual cells are labeled by cell type (top) or by experimental condition (bottom). **(E)** Gene set enrichment analysis of differentially expressed genes in pseudo-bulk (orange) or single cell (green) analysis showing the up and down- regulated pathways in post-graft B cells. **(F)** Intensity of expression of the glycolysis (left) and hypoxia (right) signatures by post-graft B cells. AGM: Aorta-gonad-mesonephros; DA: dorsal aorta; HSC: hematopoietic stem cells, FL: follicular lymphoma, E2/4: Embryonic day 2/4; NES: normalized enrichment score

Next we used scRNAseq to evaluate the effect of the engraftment procedure on fourteen FL samples. CFSE-stained cells were collected before engraftment, and CFSE+/hCD45+ cells were collected and sorted by flow cytometry 48 hours after engraftment (Supp. Figure 1A). Among the three protocols of tissue dissociation tested, collagenase combined with actinomycin D allowed the collection of a higher number of cells without inducing a transcriptomic signature associated with dissociation (van den Brink *et al*., 2017; Denisenko *et al*., 2020a) (Supp. Figure 1B). Antibody-based barcoding on pooled samples was used to minimize batch effect between cells from the same patient and scRNAseq libraries were processed using the 10X genomics technology. An average of 1455 (range 354 to 3532) and 894 (range 124 to 2776) cells per sample were found to pass the quality control of scRNAseq before and after graft respectively (Supp. Table 3). Before the graft, B cells made up 76% of all cells (range 51% to 91%) with FL cells (characterized by monotypic expression of either kappa or lambda light chain) accounting for 94% of cells for which sequencing information was available (Supp. Figure 1C). Other cells included 18% of CD4+ T lymphocytes (range 8% to 44%), including T follicular helper (6%, range 2% to 21%) and other T cells (12%, range 4% to 31%), 4% of CD8+ T lymphocytes (range 0.7% to 12%) and rare myeloid cells such as monocytes (1%, range 0.2% to 2%). The analysis of cells collected 48 hours after the graft showed a significant increase in B cells (average increase of 17%, range -6% to 27%, p<0.05) at the expense of microenvironment cells (Figure 1B). This relative enrichment in tumor cells could be partly due to a higher proportion of actively cycling tumor cells (67.9%) as compared to non-tumor cells (56.3% in S/G2M) at the time of engraftment (Figure 1C, Supp. Table 3).

Unsupervised analysis revealed that the transcriptomic diversity was primarily caused by cell type (B cells vs non-tumor cells) and experimental condition (pre vs. post-graft) (Figure 1D), which together explained 28% of the variability of the dataset by principal component analysis (Supp. Figure 1D). The intersection of both single cell and pseudo-bulk analysis identified a list of differentially expressed genes (DEG) between pre and post-graft B cells (see methods and Supp. Figure 1E) (Squair *et al*., 2021). Gene set enrichment analysis (GSEA) identified an upregulation of metabolic and pro-survival pathways (glycolysis, oxidative metabolism, mTORC1 and MYC) in post-graft samples, suggesting that the early AGM region provides a nurturing environment for FL cells (Figure 1E). Interestingly, we also identified heterogeneity among the cells collected after graft. Unsupervised clustering identified a cluster representing 13% of total cells with a high level of expression of glycolysis and hypoxia signature (Figure 1F), potentially due to engraftment in hypoxic areas of the embryos.

In conclusion, the data shows that the AGM region in avian embryos provides a favorable microenvironment for primary FL cells, which prompted us to test whether it is a suitable model to examine the therapeutic response of FL.

### FL-AVI-PDX captures clinical heterogeneity of response to RCHOP

In order to assess the ability of the FL-AVI-PDX model to capture interpatient heterogeneity of therapeutic response, we exposed the engrafted embryos to RCHOP, a commonly prescribed first-line therapy for FL patients (Salles, 2020). The maximal tolerated dose (MTD) of the RCHOP combination, used with the same stoichiometry between each drug as in clinics, was determined after a single intravenous injection of the combination (Supp. Table 4). Toxicity of each dose was assessed by measuring the growth (body surface area (BSA)) and survival of avian embryos (Jarrosson *et al*., 2021, 2023). The MTD of RCHOP (rituximab 5.13 mg/kg, cyclophosphamide 10.25 mg/kg, hydroxyadriamycine (doxorubicin) 0.69 mg/kg, vincristine 0.02 mg/kg, prednisone 0.54 mg/kg) was associated with a 86% survival rate and did not impact the embryonic BSA as compared to excipient-treated embryos (Supp. Figure 2A). The pharmacokinetic profiles of the RCHOP components were also analyzed in avian embryos blood at different time points after RCHOP intravenous injection (0.5, 3 and 8 hours). All drugs tested (rituximab, cyclophosphamide, hydroxyadriamycine, and vincristine) were detectable with a concentration peak at 0.5h followed by a slow decrease over time (Figure 2A). Compounds were detected in avian embryos at concentration 10-100 times lower than those observed in lymphoma patients (Nakagawa *et al*., 2021), yet their pharmacokinetic profiles were similar (Supp. Figure 2B). These findings suggest that the RCHOP regimen can be safely given to avian embryos, and its pharmacokinetic profile is suitable for investigating the therapeutic response of FL.

**Figure 2:**
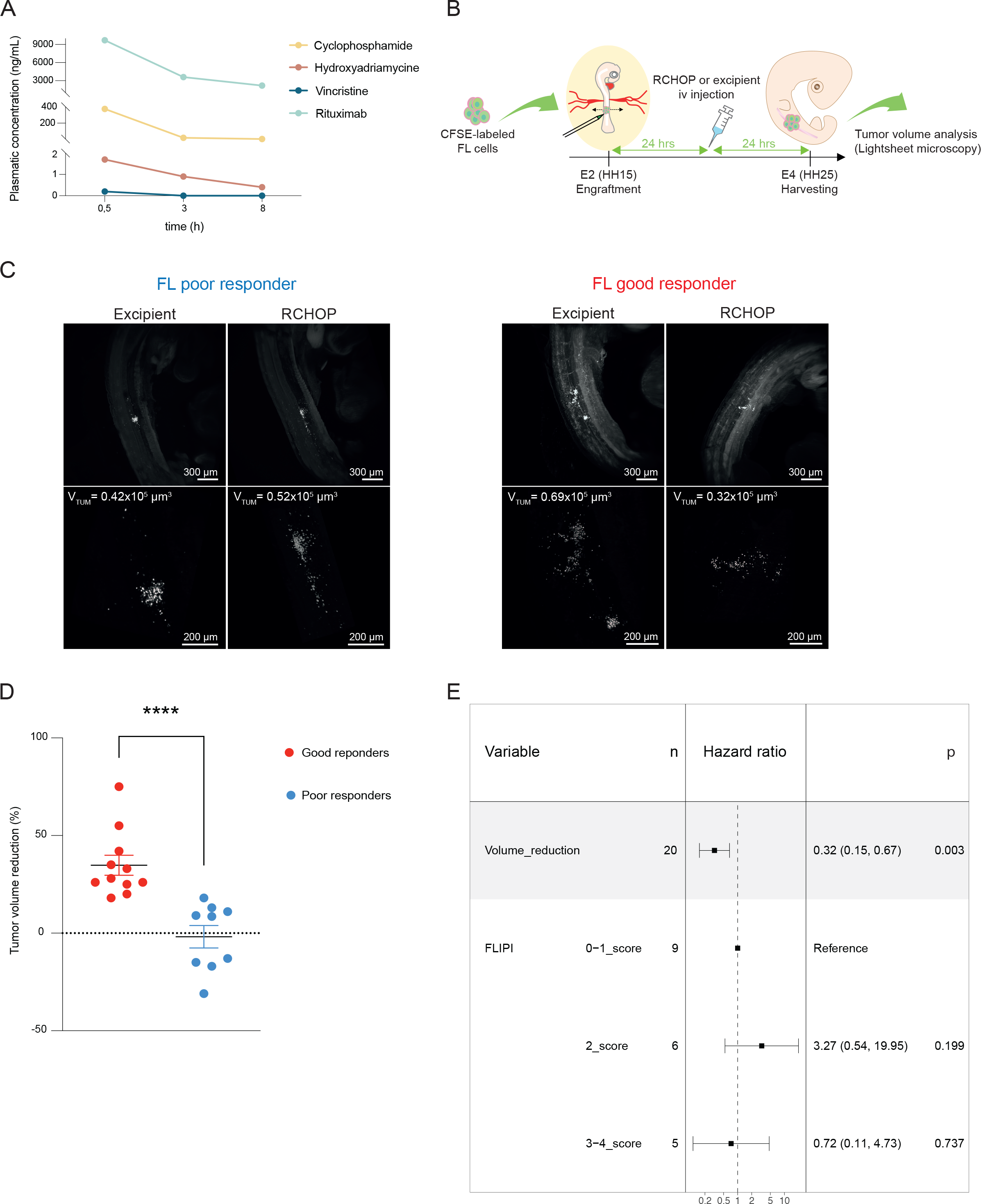
FL-AVI-PDX captures clinical heterogeneity of response to RCHOP **(A)** Plasmatic concentration of rituximab, cyclophosphamide hydroxyadriamycine and vincristine after a single intravenous injection of RCHOP MTD in avian embryos. **(B)** Schematic representation of the experimental procedure used to evaluate the effects of RCHOP on FL-AVI-PDX. **(C)** Representative example of light sheet microscopy imaging after exposure to RCHOP or excipient in a poor (left) and good responder patient (right) in whole cleared embryos (upper panels) and precise quantification of tumor volumes (lower panels). **(D)** Mean percentage of tumor volume reduction achieved with RCHOP in FL-AVI- PDX from good and poor responder patients. Error bars indicate SEM. **** p value <0.0001 (Mann-Whitney test). **(E)** Forest plot showing hazard ratios and 95% confidence intervals for progression-free survival in multivariate analysis.

We then used FL-AVI-PDX models from the twenty patients treated initially with RCHOP and rituximab maintenance, including 9 poor responders with FL progression during the first 24 months (POD24) and 11 good responders with long progression free survival (Supp. Table 1). Twenty-four hours after engraftment, FL-bearing avian embryos were randomly treated by intravenous injection RCHOP or excipient. The embryos were harvested after an additional 24 hours (HH25) for measurement of tumor volumes based on CFSE fluorescence using light sheet microscopy (Figure 2B). RCHOP consistently reduced the mean tumor volume in FL- AVI-PDX from the good responder patients but not in FL-AVI-PDX from poor responder patients (Figure 2C). Accordingly, the mean tumor volume reduction from all the samples was significantly higher in FL-AVI-PDX from good responders compared to poor responders (mean tumor volume reduction 35% vs. -2%, p=0.0001) (Figure 2D, Supp. Table 2). To further explore this result beyond the binary categorization of clinical response, we evaluated whether tumor volume reduction could predict progression free survival using a Cox model (Supp. Figure 2C). Multivariate analysis showed that tumor volume reduction, and not FL International Prognostic Index (FLIPI), is an independent prognostic predictor in this cohort (hazard ratio 0.32, range 0.15-0.67, p=0.003) (Figure 2E). Altogether, these data show that the FL-AVI-PDX heterogeneous response to RCHOP is correlated to clinical outcome.

### FL-AVI-PDX reveals a transcriptomic signature of response to RCHOP in primary FL cells

The knowledge about the mechanisms of action of chemotherapies on FL cells is mostly derived from cancer cell lines, and no data have been generated on primary cells so far. Therefore, we further used the FL-AVI-PDX model to explore the mechanisms of response of primary FL cells to RCHOP by using scRNAseq. We compared B cells from fourteen FL- AVI-PDX collected after graft and exposed to either RCHOP or vehicle, by combining both single cell and pseudo-bulk strategies in order to mitigate the batch sample effect observed in single cell analysis. A list of 21 genes upregulated after RCHOP exposure was identified by both methods (Figure 3A, Supp. Table 5). Among these genes are key effector of p53 response and/or apoptosis such as *DDB2, PVT1, BAX* and *BBC3* (*PUMA*). We confirmed the induction of some of these genes (*BAX, RPS27L*, *RPS19* but not *SRSF3*) after RCHOP exposure by quantitative PCR in a germinal center derived lymphoma cell line (SUDHL-4) (Supp. Figure 3A). We then treated this line with each drug separately, to analyze their contribution to the transcriptomic modifications identified in response to RCHOP. Except for *RPS27L* whose induction was only observed with hydroxyadriamycine, the induction of the other genes was not observed with any single drug, suggesting that complex synergies drive the observed transcriptomic modifications (Supp. Figure 3B).

**Figure 3:**
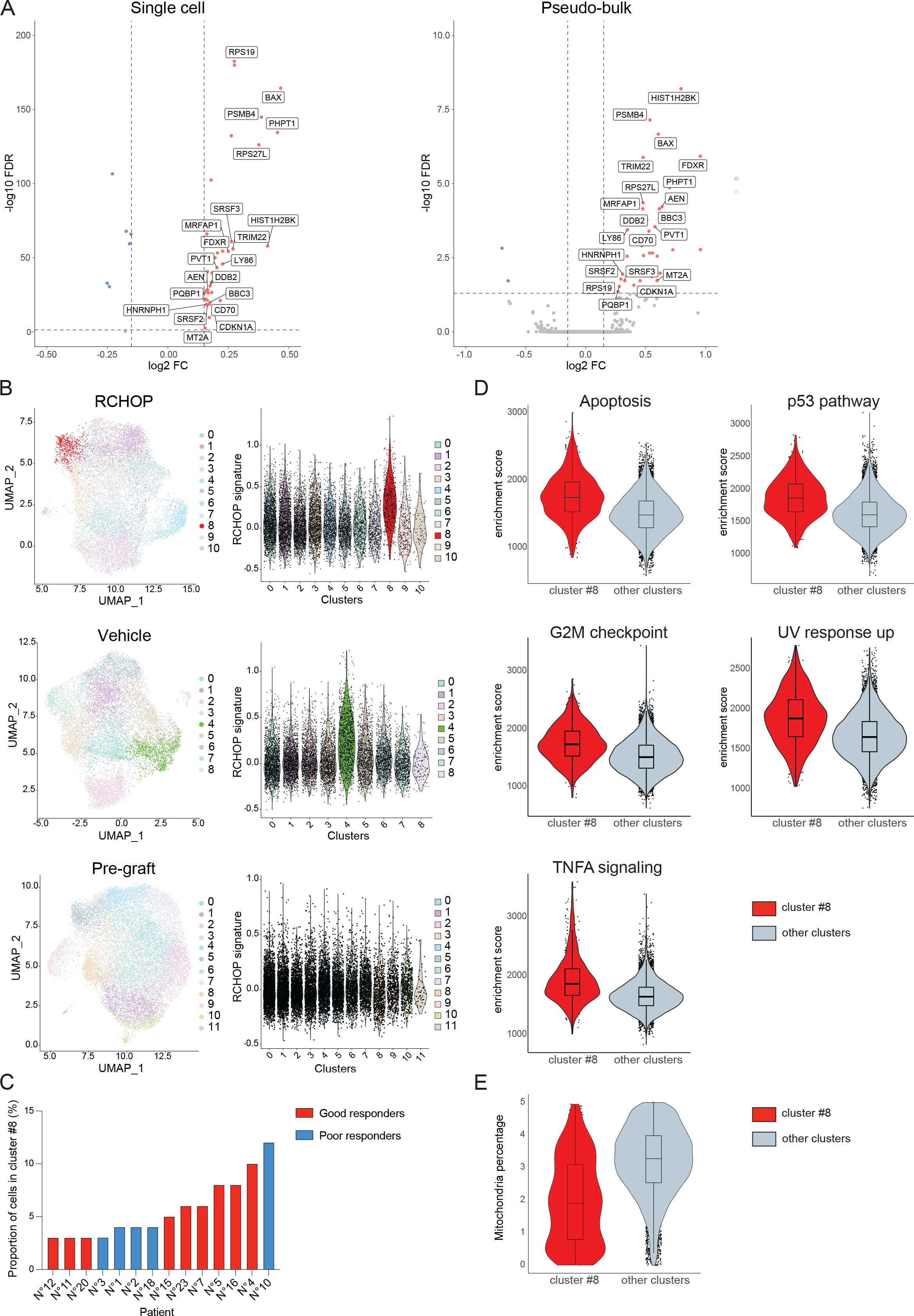
FL-AVI-PDX reveals a transcriptomic signature of response to RCHOP in primary FL cells **(A)** Volcano plot showing the differentially expressed genes by B cells exposed to RCHOP as determined by single cell (left panel) and pseudo bulk method (right panel). The 21 up- regulated genes identified by both methods are labeled in red. **(B)** Uniform Manifold Approximation and Projection (UMAP) representation of the scRNAseq analysis of B cells exposed *in ovo* to RCHOP (upper panel), to vehicle (central panel) or before graft (lower panel) and intensity of the expression of the RCHOP signature in each cluster for each corresponding condition. **(C)** Percentage of cells belonging to cluster #8 for each sample (n=14, including 9 good responders (red) and 5 poor responders (blue)) **(D)** GSEA analysis of cells belonging to cluster #8 as compared to cells from all other clusters in RCHOP-exposed cells (FDR < 0.05). (**E**) Violin plot representation of the percentage of genes encoded by the mitochondrial genome in cells belonging to cluster #8 as compared to cells from all other clusters in RCHOP-exposed cells.

Intratumoral heterogeneity is recognized as an important factor driving resistance to cancer treatment, which is especially important in FL where relapses occur invariably. To evaluate the heterogeneity of response to RCHOP among cancer cells, we assessed the level of expression of the RCHOP signature in the different B cells clusters identified by unsupervised clustering strategies. Interestingly, we found that a specific cluster (cluster #8) strongly expressed the RCHOP signature. We also identified a cluster (cluster #4) overexpressing this signature in samples stressed only by the graft procedure, albeit with lower magnitude as compared to RCHOP exposed samples, and no such cluster was found in pregraft samples. (Figure 3B). Cluster #8 was present in each patient’s sample, representing 3-12% of the cells, and its relative size was not different between samples from good or poor responders (Figure 3C, Supp. Figure 3C). Further analysis using GSEA showed that this cluster was significantly enriched in transcriptomic signatures of apoptosis, p53 activation, cell cycle checkpoint, UV response and TNF alpha signaling (Figure 3D). Unexpectedly, expression of mitochondrial genes, which is commonly used as a proxy of apoptotic process, was significantly lower in this cluster (Figure 3E).

Altogether, our findings uncover a transcriptomic signature of response to RCHOP in primary FL cells and highlight heterogeneity in the level of induction of this signature among primary FL cells. To further validate the FL-AVI-PDX model, we investigated whether we could identify new treatment approaches in FL based on the transcriptomic signature of response to RCHOP.

### *BAX* is a targetable effector of RCHOP in FL

*BAX* was found to be overexpressed after RCHOP exposure in primary cells, and especially in cluster #8 cells, as well as in the SUDHL-4 cell line (Figure 4A, Supp. Figure 3A). *BAX* interacts with BH3-only proteins such as BAD, BID, or BIM, and forms homo- or hetero-oligomers (with BAK) in the outer mitochondrial membrane, causing cytochrome c release and apoptotic cell death through the process of mitochondrial outer membrane permeabilization (MOMP). Its activation is inhibited by pro-survival members of the BCL2 family such as BCL2 itself, which is overexpressed in FL due to the t(14;18) translocation (Huet, Sujobert and Salles, 2018).

**Figure 4:**
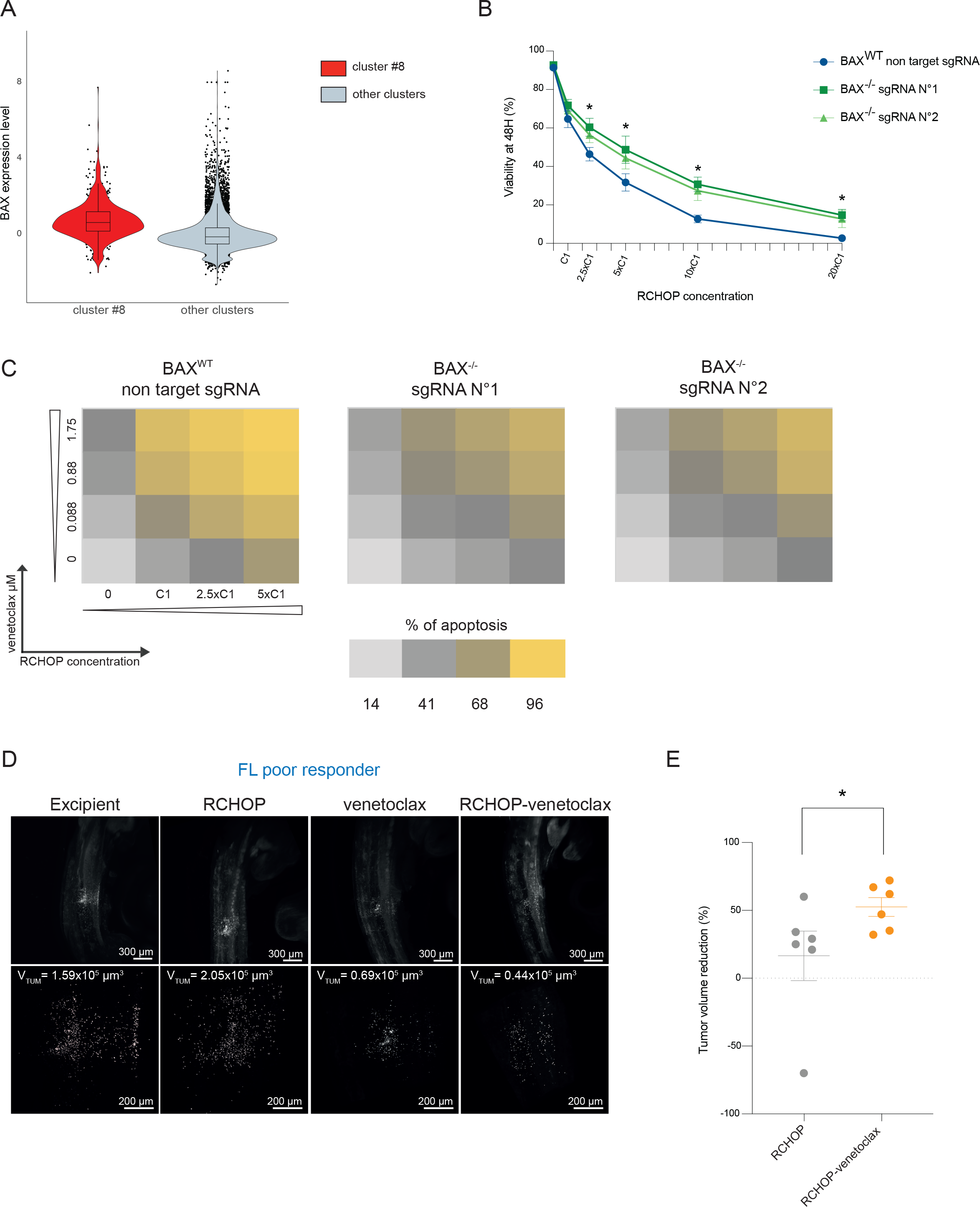
*BAX* is a targetable effector of RCHOP in FL **(A)** Violin plot representation of *BAX* level expression in cells belonging to cluster #8 as compared to cells from all other clusters in RCHOP-exposed cells. **(B)** Proportion of viable SUDHL-4 cells (*BAX*^-/-^ with two different sgRNA targeting *BAX* or *BAX*^WT^ with a non-target sgRNA) after 48h exposure to increasing doses of RCHOP. Error bars indicate SD. *p Val<0.05 (Wilcoxon test). **(C)** Proportion of apoptotic SUDHL-4 cells (*BAX*^-/-^ with two different sgRNA targeting *BAX* or *BAX*^WT^ with a non-target sgRNA) after a 48h-exposure to increased doses of RCHOP and venetoclax combinations. **(D)** Example of light sheet imaging for one embryo of each condition in whole cleared embryos (upper panels) and precise quantification of tumor volumes for analysis (lower panels) for a patient with poor response to RCHOP. **(E)** Mean percentage of tumor volume reduction observed after RCHOP or RCHOP-venetoclax treatment of FL-AVI-PDX (n=6). Error bars indicate SEM, * p<0.05 (Mann-Whitney test).

First, we examined the effect of *BAX* inactivation on RCHOP-induced apoptosis. Using CRISPR-Cas9, we deleted *BAX* in the SUDHL-4 cell line with two different sgRNA (Supp. Figure 4A). The *BAX*^-/-^ cell lines were less sensitive to RCHOP compared to the wild type cells expressing a non-targeting sgRNA (Figure 4B). We hypothesized that increasing the level of *BAX* or its activation might, on the contrary, further enhance the effects of RCHOP. To test this hypothesis we used venetoclax, an FDA-approved BCL2 inhibitor that indirectly increases *BAX* activity (Delbridge *et al*., 2016; Blombery *et al*., 2022) (Supp. Figure 4B). As expected, *BAX*^-/-^ cells were resistant to venetoclax (Supp. Figure 4C). The combination of RCHOP and venetoclax resulted in an additive effect on cell death induction in the SUDHL-4 lymphoma cell line, in a *BAX*-dependent manner (Figure 4C).

To further examine the potential benefits of targeting *BAX* to enhance the efficacy of RCHOP, we compared the RCHOP-venetoclax combination to RCHOP or venetoclax alone using the FL-AVI-PDX model. We first established the MTD of venetoclax when used alone or in combination with RCHOP. The MTD of intravenous venetoclax as a single agent was 172 mg/kg, but it decreased to 57 mg/kg when combined with RCHOP (Supp. Figure 4D, Supp. Figure 4E and Supp. Table 6). We used this dose to assess the efficacy of the RCHOP- venetoclax combination in six primary FL samples from good (n=4) or poor responders (n=2) to RCHOP (Supp. Table 1). As expected, the FL-AVI-PDX established from poor responder patients did not respond to RCHOP alone, but a pronounced response was observed with the addition of venetoclax (Figure 4D). The better efficacy of the RCHOP-venetoclax was confirmed in the whole cohort (mean tumor volume reduction 52.5% vs. 16.5% p<0.05) (Figure 4E). These findings demonstrate that the FL-AVI-PDX model can be used to identify and validate combination therapies.

## Discussion

For decades, researchers have focused on cell lines as models for cancer. However, with the progressive recognition of the ecological complexity of cancer (Dujon *et al*., 2021), the development of more relevant models has become necessary. Patient-derived xenografts in immunocompromised mice have been a major breakthrough, allowing for direct evaluation of drug therapies in a more physiological setting (Townsend *et al*., 2016). However, this strategy has limitations for B cell lymphomas, particularly for FL, which grow only occasionally in these models, and have a long latency of development (Zhang *et al*., 2017). Here we report experimental data supporting the use of avian embryos as an alternative recipient for FL PDX. This model allowed for successful engraftment of all samples tested, with a large number of replicates, providing robust statistical power to test therapeutic strategies. Using RCHOP as a proof of concept, we observed a clear correlation between response to RCHOP *in ovo* and clinical outcome, which demonstrates that clinically relevant information can be derived from functional testing of FL in this model. As almost all FL patients achieve complete clinical response after 6 courses of RCHOP, the observation of a significant difference of response to RCHOP *in ovo* was unexpected. We hypothesize that this differential effect on bulk tumor reduction is due to the low drug concentrations achieved in avian embryos, which may reveal subtle differences in chemosensitivity. Other limitations of the model are the absence of complement proteins in avian embryos at the early stages of development, which might limit the activity of rituximab (Di Gaetano *et al*., 2003), and short experiment duration. Further research is necessary to determine if longer duration of graft could provide a deeper understanding of inter-patient variability. Another potential development of this model as a predictive biomarker would require prospective evaluation in large patient cohorts.

The FL-AVI-PDX model also allows to collect back cells from the grafted tumors, which provides a unique opportunity to examine the effects of treatments on primary tumor cells. We describe here a list of 21 genes that were upregulated after RCHOP exposure, including genes related to the known mechanisms of action of chemotherapy. Notably, not all of these genes were activated after RCHOP treatment of a cell line, emphasizing the value of using FL PDX models to fill in the knowledge gaps. As a proof of concept of the usefulness of this model, we performed functional validation of the role of *BAX* in RCHOP-induced apoptosis. This led us to investigate the potential benefits of combining venetoclax with RCHOP, which was found to be additive in both cell lines and primary FL *in ovo*, including in samples from poor responder patients. Of note, the RCHOP-venetoclax combination has been already evaluated in the CAVALLI phase 1b study which included 10 FL patients, and showed acceptable safety profile and a complete remission rate of 70% (Zelenetz *et al*., 2019). The observed heterogeneity in the treatment response *in ovo* suggests that the results at the cohort level might underestimate the potential benefit of this combination for subgroups of patients such as those predicted to poorly respond to RCHOP.

In addition to its use as a platform for preclinical development in FL treatments, the FL-AVI-PDX model revealed intratumoral heterogeneity in response to RCHOP. In all samples we identified a cluster of cells (cluster #8) with a high expression of the RCHOP signature. This signature reveals the activation of stress response (such as the p53 pathway), which is also heterogeneously expressed in engrafted samples not exposed to chemotherapy, albeit to a lesser extent. Intriguingly, the expression of mitochondrial genes, which are usually considered as a proxy of the apoptotic process in single cell RNA-seq studies, was lower in cells belonging to cluster #8. This raises the hypothesis that these cells may be able to withstand strong proapoptotic gene expression due to their low mitochondrial content (Márquez-Jurado *et al*., 2018). Further study through the isolation of these cells is needed to confirm this hypothesis and assess their role in relapse after immunochemotherapy.

## Materials and methods

### Patient Samples

Diagnostic biopsy samples from FL patients treated with RCHOP followed by rituximab maintenance were mechanically dissociated and cryopreserved in 10% fetal calf serum (FCS) and dimethylsulfoxyde (DMSO). All patients gave their written consent for the use of their samples in research protocol. Electronic health records were reviewed to determine the main characteristics of the patients at diagnosis and their outcome (Supp. Table 1). Progression free survival (PFS) was defined as the delay between treatment initiation and relapse, progression or death.

### *In ovo* xenograft of Follicular Lymphoma samples (FL-AVI-PDX)

The AVI-PDX procedure is a registered and patented trademark of OncoFactory SAS, an ERBC company.

#### Avian embryos

Embryonated chicken eggs were obtained from a local supplier (Élevage Avicole du Grand Buisson, Saint Maurice de Dargoire, France). Laying Hen’s sanitary status was regularly checked by the supplier according to French laws. Eggs were housed in an incubator at 18°C until further use. They were then incubated at 38.5°C in a humidified incubator until the desired developmental stage. In all experiments, embryos were randomized in each experimental group and were harvested at embryonic day 4 (4 days post-fertilization).

#### Graft procedure

After thawing, FL cells were labeled with an 8μM Carboxy Fluorescin Succinimidyl Ester (CFSE) solution (Life technologies Carlsbad, California) and a fraction of CFSE-labeled cells was frozen at this step (pre-graft experimental condition). Stage HH14 avian embryos were grafted with 1000-2000 cells in the presumptive AGM region, with a glass capillary connected to a pneumatic pico-pump (PV820, World precision instruments, Sarasota, Florida) under a fluorescent stereomicroscope. Grafted eggs were incubated at 38.5°C in a humidified incubator until HH25 stage, as already published (Delloye-Bourgeois *et al*., 2017; Jarrosson *et al*., 2021).

#### Drug administration and determination of drug maximum tolerated dose (MTD) in avian embryos

RCHOP and venetoclax compounds were obtained from the pharmacy of Centre Léon Bérard hospital (rituximab, cyclophosphamide, doxorubicin) or purchased from Selleckchem (vincristin, prednisone and venetoclax). All compounds were diluted in DMSO 95%, NaCl 0.5% and 0.5% Tween 80, used as an excipient solution for *in vivo* experiments.

For the determination of drugs maximum tolerated dose, increasing doses were injected intravenously into the chorioallantoic vessels, with a single administration in HH20 embryos. Twenty-four hours after injection, avian embryos were harvested, weighed (Sartorius Quintix35-1S) and measured along the rostro-caudal axis using the Leica LASX image analyses software. The Body Surface Area (BSA) was calculated using Dubois & Dubois formula: BSA (m^2^) 0.20247 x height (m)^0.725^ x weight (kg)^0.425^. The morphology / anatomy of each embryo was systematically analyzed to check their correct stage-related development. The criteria observed were the survival (heart beating), the craniofacial morphology (presence of each cerebral compartment and eyes), the presence of four limb buds, the cardiac morphology, and the anatomy of embryonic annexes such as the allantois. An embryonic death rate over 70% or the observation of developmental anomalies in more than 50% of embryos was considered to be indicative of drug toxicity.

#### Pharmakokinetics in avian embryos

RCHOP was injected into the bloodstream of avian embryo at embryonic day 3 (E3). Then, blood was collected from extraembryonic vein in a series of avian embryo 0.5, 3, and 8 h after RCHOP administration. A total of 15µL of blood plasma was collected for each condition, in duplicate at each time points. 10 µL of internal standards solution, 600 µL of phosphate buffer (50 mM, pH: 7.2), and 3 mL of a mixture of methanol/chloroform (25/75) were added successively to the volume of plasma. The sample was mixed vigorously for 10 min and then centrifuged for 5 min at 4000 g at room temperature. The organic layer was transferred to a glass tube and evaporated at 45°C under a gentle stream of nitrogen. The residue was reconstituted in 100 µL of mobile phase and 10 µL were injected into the apparatus. Analysis was performed on a liquid chromatography coupled with high resolution mass spectrometer (LC-MS/HRMS - Q-Exactive Plus Orbitrap; ThermoFisher Scientific™). The electrospray source operated in positive mode. The separation of the compounds was carried out with a Hypersil Gold® column (ThermoFisher Scientific) and a stepwise gradient program. The sample preparation was based on liquid/liquid extraction and cyclophosphamide [^2^H8], daunorubicin and vindesine were used as internal standards for cyclophosphamide, doxorubicin and vincristine, respectively.

The quantification of rituximab was performed with a LC-MS/HRMS method previously described (Millet *et al*., 2021).

#### Tissue clearing and whole mount Selective Plan Imaging Microscopy (SPIM) imaging

PFA-fixed HH25 embryos were cleared using an adapted Ethyl-Cinnamate protocol (Jarrosson *et al*., 2021). Briefly, tissues were dehydrated in ethanol successive baths and finally cleared in Ethyl Cinnamate (Sigma, 112372). Cleared samples were imaged using the UltraMicroscope SPIM (Miltenvi Biotec). 3D-images were built using Imaris™ software. Volumetric analysis was performed using Imaris^TM^ ^“^Surface” module adjusted on CFSE fluorescence. Statistical treatment of the data was performed with Prism 9.0e (GraphPad). Both normality (d’Agostino & Pearson test) and variance homoscedasticity (*F* test) were checked. In cases where experimental groups did not pass normality, non-parametric tests were used. In cases where experimental groups passed normality tests but had significantly difference variances, corrections were applied (Welch’s correction *t*-test). All statistical tests were two-sided. The exact test and p-values are mentioned in the figure legends and supplementary tables.

#### Flow cytometry and cell sorting

FL tumors formed in avian embryos treated with the excipient or RCHOP were microdissected out 48h after grafting procedure using a stereomicroscope. Microdissected tumors were dissociated in Hank’s Balanced Salt Solution (HBSS) with 156 units/ml of type IV collagenase, 15mM Actinomycin D and 50 units/ml of DNase I for 10 min at 37°C, and then incubated with 5mg/ml trypsin for 1 min at 37°C under gentle mixing. Actinomycin D was added at all steps of the protocol to limit the impact of dissociation on transcriptomic data. Ultimately, Dulbecco’s Modified Eagle’s Medium (DMEM) supplemented with 10% Foetal Bovine Serum (FBS) and 3mM Actinomycin D were added to inhibit enzymatic activity. Non-dissociated tissue was removed by filtration through 0.4mM nylon cell stainer (BD Falcon). Dissociated cells were frozen down in 10%DMSO - 20% FCS - 70% RPMI using cell cooler and stored at -80°C prior to cell sorting. Frozen cell suspensions were thawed at 37°C in RPMI with 20% FCS, washed in phosphate buffer saline (PBS) and resuspended between 50-150x10^6^cells/mL in FACS buffer (PBS with 5% FCS and 2mM EDTA). Cells were first blocked with 2% mouse serum (eBioscience San Diego, California) and Human Fc Block (Biolegend San Diego, California) for 10 minutes on ice. Cells were then incubated with a pre-mixed antibody solution of human CD45 APC-Cy7 (Becton,

Dickinson and Company Franklin Lakes, New Jersey) and hashtag barcoded antibodies (Biolegend) specific for each experimental condition (pre-graft, RCHOP and vehicle) for 30 minutes on ice. After washing with FACS buffer, cells were resuspended in the same buffer at 1x10^7^/mL. DAPI (Sigma-Aldrich St Louis, Missouri, ref #D9542) was added immediately before the cell sorting procedure using a FACS Aria III cytometer (Becton, Dickinson and Company). The gating strategy is described in Supplementary Figure 1A.

### Single cell RNA-seq

#### Wet lab

Sorted cells were immediately loaded into the Chromium Controller (10X genomics, Pleasanton, California) and hashtag libraries were prepared according to the manufacturer’s instructions (Next GEM Single Cell V(D)J V1.1 with feature barcoding technology workflow). The V(D)J library was amplified from 5-20 ng of cDNA with two PCR of 6 and 8 cycles for the heavy and light chains respectively with primers recommended by 10X Genomics. After solid phase reversible immobilization (SPRI) cleanup, 800 pg of amplified sequences was used with Nextera XT reagents (Illumina San Diego, California) to construct the final BCR library by PCR using i5 forward and Nextera XT i7 reverse primers. The libraries were quantified using the Library Quantification Sample Kit Kapa (Illumina-Uni San Diego, California). Gene expression, hashtag and V(D)J libraries were pooled in a 8/1/1 ratio, mixed with 1% PhiX and paired-end (2x150pb) sequenced (375M reads) on a HiseqX platform (Macrogen Seoul, South Korea) (Supp. Table 5).

#### Creation of the Seurat object and normalization

BCL files were converted to FASTQ using CellRanger mkfastq (R1 : 26 base pairs, R2 : 150 base pairs). Raw FASTQ from gene expression libraries were aligned on the GRCh38 human genome and processed using CellRanger Count (10x Genomics Cell Ranger 4.0.0). HTO barcodes were retrieved using CITE-seq-Count and normalized for each cell using the centered log ratio (CLR) transformation implemented in the Seurat NormalizeData function (Hao *et al*., 2021). Cells were demultiplexed using the Seurat HTODemux function, and barcodes assigned as doublets or negative were excluded from further analysis. Quality controls were used to exclude cells with a number of detected genes below 200 and above 2500 and cells with more than 5% of transcripts encoded by the mitochondrial genome. The resulting filtered UMI count matrices were log-normalized using Seurat NormalizeData with a scale factor of 10.000 and a scaling step was performed using Seurat ScaleData with all genes from the matrices as features.

#### Metadata creation

Cell phenotype annotations were identified using singleR and Celldex R packages against cell markers from the MonacoImmuneData database (Monaco *et al*., 2019).

Cell cycle analysis was performed using Seurat CellCycleScoring. Cells were assigned to G2/M or S phase based on their highest expression score, while cells expressing neither were assigned to G0/G1 phase.

B cell immunoglobulin phenotype reconstruction was performed with CellRanger VDJ (10x genomics Cell Ranger 4.0.0) using FASTQ files from the VDJ library and the GRCh38 reference genome. BCR contigs contained in ’filtered_contig_annotations.csv’ were manipulated in R to obtain immunoglobulin light and heavy chains for each cell.

The Seurat function AddModuleScore was used to calculate the average expression levels of the glycolysis signature (Molecular Signature Database (Liberzon *et al*., 2011) (MSigDB), n=186 genes), the hypoxia signature (MSigDB, n=182 genes), the dissociation signature (published by Denisenko et al., n=427 genes (Denisenko *et al*., 2020b)) and the RCHOP signature identified in the present study (n=21 genes). All calculations were made from the “RNA” assay of the Seurat objects.

#### Data integration

We integrated the sample-specific objects using the Seurat v4. Each filtered UMI count matrix was normalized a second time with Seurat SCTransform (vst.flavor = “v2”) and dimension reduction was performed with RunPCA (npcs=5). Integration was performed using SelectIntegrationFeatures (nfeatures = 3000), FindIntegrationAnchors (dims= 1:20) and IntegrateData (dims= 1:20). The resulting integrated object was centered, reduced with RunPCA and embedded in two dimensions with RunUMAP (umap.method= “umap-learn”), excluding BCR- and TCR-encoding genes (Denisenko *et al*., 2020a; Sundell *et al*., 2022) from the lists of variable genes (regex: IG[HKL][VDJ] |IGHG[1-4]| IGH[MDE] |IGKC |IGLL |IGLC[1-7] |IGHA[1-2] |TR[ABGD][CV]) determined by the Seurat function VariableFeatures. These genes were also removed from the matrix count for further analyses.

#### Pseudo-bulk aggregation

Pseudo-bulk matrices were computed using the aggregation method from the HBC training tutorial (Khetani, 2020). Starting from the raw matrices merged from all single cell samples, poor quality cells were considered as outliers based on median-absolute-deviation (MAD) method (scater package, isOutlier function, nmads = 2). B cells from the same sample and the same experimental condition (pre-graft, post-graft-vehicle, and post-graft-RCHOP) were aggregated. As a final quality control, the pseudo-bulk matrix computed using the filterByExpr function from the edgeR package was applied once (Chen, Lun and Smyth, 2016). The remaining pseudo-bulk matrix was lognormalized using the trimmed mean of M (TMM) method(Robinson and Oshlack, 2010; Denisenko *et al*., 2020a) performed by the edgeR calcNormFactors function.

#### Identification of differentially expressed genes

Single-cell differential gene expression lists were calculated using Seurat FindMarkers (test.use= “bimod”, assay= “RNA”) from integrated matrices. The LogFC threshold was set to 0.15 for the RCHOP vs. vehicle (post-graft) assay. Only B cells were retained for testing.

The EdgeR glmTreat approach was used from pseudo-bulk matrices to test for differential expression with a fold change threshold of 1.11 and a false discovery rate (FDR) below 0.05 to be comparable to the logFC threshold from the single cell differential expression test.

The differentially expressed genes retained were those identified by both approaches.

#### Gene Set Enrichment Analysis (GSEA)

At the single cell level, GSEA was performed using the enrichIt function of the escape R package (Borcherding *et al*., 2021). The minimum number of genes required to perform the enrichment calculation was set to 5 and only pathways with FDR< 0.05 were kept. At the pseudo-bulk level, fast gene set enrichment analysis (fGSEA) was performed using the fgsea function of the fgsea R package, applying the hallmark collection (50 gene sets, h.all.v7.5.1.symbols.gmt) from MSigDB. The log2(FC) value of each gene, previously calculated by the edgeR package from pseudo-bulk processing, was used as the ranking metric input. All pathways smaller than a size of 5 gene set to test was excluded.

### In vitro experiments

#### Cell culture

The SUDHL-4 cell line (CVCL_0539, male derived) with constitutive expression of the Cas9 protein was grown in Roswell Park Memorial Institute medium 1640 GlutaMAX Supplement (ThermoFisher Scientific Waltham, Massachusetts) supplemented with 20% FCS, 1% penicillin G/streptomycin (5000U/mL) (ThermoFisher Scientific). Cells were passed in fresh medium every 3 days and grawn in incubators at 37°C in a 5% CO2 atmosphere.

RCHOP compounds were purchased from the hospital pharmacy facilities of Hôpital Lyon Sud (Lyon, France) and venetoclax (ABT-199) was purchased from Selleck Chemicals (Houston, Texas). For drug sensitivity assays, cells were seeded at 4x10^5^ cells/mL in 96-well flat bottom culture plates. The RCHOP concentrations used were as follows in a final volume of 200µL/well:

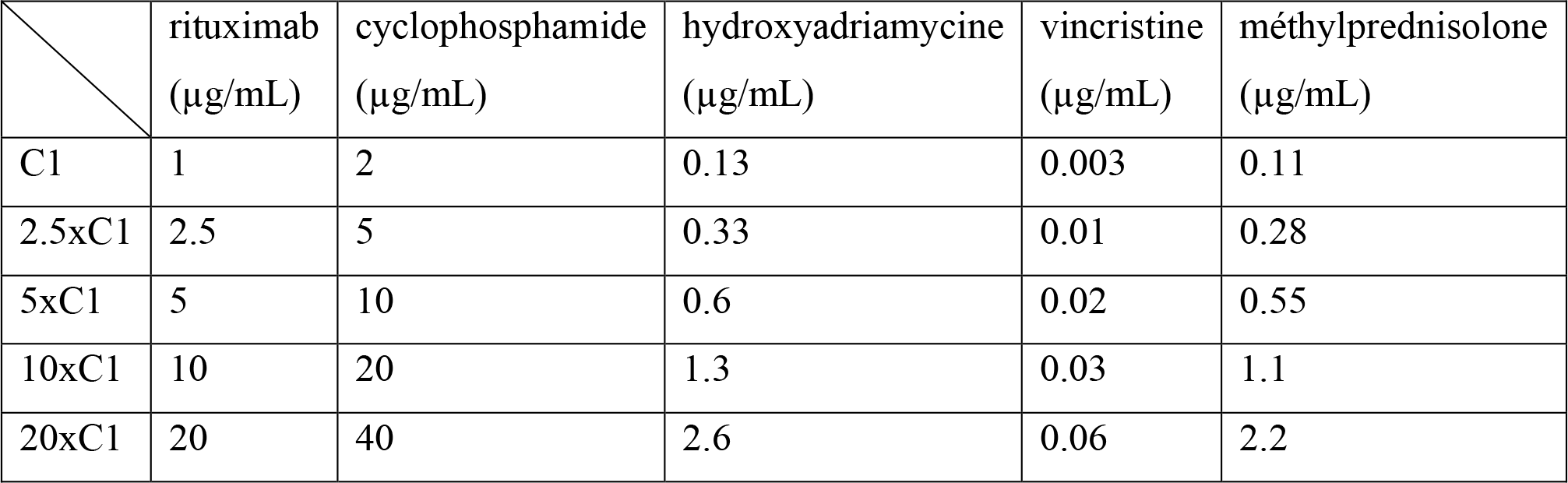

|  | rituximab<br>(µg/mL) | cyclophosphamide<br>(µg/mL) | hydroxyadriamycine<br>(µg/mL) | vincristine<br>(µg/mL) | méthylprednisolone<br>(µg/mL) |
| --- | --- | --- | --- | --- | --- |
| C1 | 1 | 2 | 0.13 | 0.003 | 0.11 |
| 2.5xC1 | 2.5 | 5 | 0.33 | 0.01 | 0.28 |
| 5xC1 | 5 | 10 | 0.6 | 0.02 | 0.55 |
| 10xC1 | 10 | 20 | 1.3 | 0.03 | 1.1 |
| 20xC1 | 20 | 40 | 2.6 | 0.06 | 2.2 |

Viability was assessed 48h later after labeling cells with Annexin V APC (TONBObiosciences San Diego, California) on the ATTUNE NxT cytometer instrument (Life technologies Carlsbad, California) and analyses were performed using FlowJo software (v10.8.1, Becton, Dickinson and Company Franklin Lakes, New Jersey). Each experiment was performed in biological triplicates.

#### Generation of BAK^-/-^ SUDHL-4 cell line

SUDHL-4 cell line was infected using pLKO.1-puro-GFP U6 sgRNA (Addgene #50920) containing BAX RNA guides or non-targeting RNA guide:

BAX-KO-F sgRNA N°1: ACCGTCGGAAAAAGACCTCTCGGG BAX-KO-F sgRNA N°2: ACCGAGTAGAAAAGGGCGACAACC Non-Targeting-F: ACCGGCAGTACTACTGAGTTTTTC Cells were selected using puromycin (3µg/mL) during 7 days.

#### Immunoblotting

Prior to immunoblotting, cells were lysed using cOmpleteTM, Mini, EDTA-free protease inhibitor cocktail (Roche Basel, Switzerland) with the addition of phosphatase inhibitor cocktails (Sigma Aldrich). Proteins were assayed using BC Assay Protein Assay Kit (Interchim Montluçon, France) and 30µg was used for immunoblotting experiment. Lysates were separated by SDS-PAGE and transferred to nitrocellulose membrane using iBlot2 device (ThermoFisher Scientific Waltham, Massachusetts). Rabbit anti-BAX (Cell Signaling Danvers, Massachusetts, ref #2774), rabbit anti-BAK (Cell Signaling, ref #3814), and mouse anti-β-actin (Cell Signaling, ref #3700) were used as primary antibodies and were detected with anti-rabbit or anti mous IgG HRP-conjugated secondary antibodies (Cell Signaling, ref #7074 and #7076). Proteins were detected by chemiluminescence using Immobilon Forte Western HRP substrate (Merk Rahway, new Jersey) on the Chemiluminescence Western Blot Imaging System Azure C300 instrument (Azure Biosystem Dublin, California).

#### RT-qPCR

Relative expression analysis of BAX, RPS27L, SRSF3 and RPS19 was performed on SUDHL-4 cell line after overnight exposure to RCHOP or vehicle as described above. The dose used for incubation with RCHOP was Dose 4 and incubation with individual drugs was performed with the same final concentrations of each compound.

Total RNA extraction was performed using the RNeasy Mini Kit (Qiagen Hilden, Dutch) according to the manufacturer’s instructions. Each extraction was performed after counting cell viability with trypan blue to ensure at least 70% viability.

Reverse transcriptions were performed from 1 µg of RNA using M-MLV reverse transcriptase (Invitrogen Waltham, Massachusetts). After 1/100 dilution of cDNA, each transcript was amplified using QuantiTect Primer Assays QT00031192, QT00204750, QT00052829, QT00058142, and QT01192646 (Qiagen) for the detection of BAX, RPS27L, SRSF3, RPS19, and GAPDH, respectively. Amplification was performed using QuantiTect SYBR Green PCR Master Mix (Qiagen), and all RT-qPCR reactions were performed on an AriaMx Real-Time PCR System instrument (Agilent Technologies Santa Clara, California). Thermal cycling conditions were as follows:

SRSF3 and RPS19 genes: 95°C for 15min and 40 cycles of (95°C for 30sec; 55°C for 1min) BAX gene: 95°C for 15min and 40 cycles of (95°C for 30sec; 55°C for 45sec) RPS27L gene: 95°C for 15min and 40 cycles of (95°C for 30sec; 55°C for 30sec) GAPDH gene: 95°C for 15min and 40 cycles of (95°C for 30sec; 55°C for 1min; 72°C for 30sec) Relative quantification of gene expression was determined using the 2-ΔΔCT method with GAPDH as the reference gene. Each data point was performed in experimental and biological triplicates.

### Kaplan-Meier and multivariate analyses

The progression free survival (PFS) curve was made from the survfit function (survival package v3.5.3) and multivariate Cox hazard regression model was used to evaluate the relationship between PFS and FLIPI or tumor volume reduction (coxph function). Forest plot was used as a visualization of the Cox model using forestmodel package (v 0.6.2). p-values of less than 0.05 were considered statistically significant.

#### Data Sharing Statement

The single-cell RNA-seq data generated during the current study is available in the Gene Expression Omnibus database under accession code GSE231523.

## Acknowledgements

The authors thank the Institut Carnot Calym for having provoked the scientific collaboration and provided access to the CeVi_Collection. The authors also thank the Centre de Ressources Biologiques CRB-Sud from Hospices Civils de Lyon, Lyon Sud hospital pharmacy department for having provided RCHOP compounds. We thank Yann Guillermin for his help in data collection. We thank Sabrina Baaklini, Pierre Milpied, Bertrand Nadel, Jerome Tamburini, and Bruno Tesson for helpful scientific discussions. We acknowledge the contribution of SFR Biosciences (UAR3444/CNRS, US8/Inserm, ENS de Lyon, UCBL) lentivectors production facility (Gerland) and cytometry facility (Lyon Sud). We thank ChatGPT for English editing.

## Authorship Contributions

M.Z., L.G., V.C., C.D-B., and P.S. conceptualized the study and acquired fundings; M.Z., B.L., C.C., L.J., R.T., M.L., A.V., and J.G., performed the investigations; M.Z., B.L., C.C., L.J., R.T., E.J., M.L., V.C., C.D-B., and P.S. designed the methodology; A.T-G., E.B., G.S., S.H., P.S. provided resources; M.Z., R.T., V.C., C.D-B., and P.S. supervised the study; M.Z and P.S. wrote the original draft.

