## Supplemental Figures and Tables for "Functional precision oncology for follicular lymphoma with patient-derived xenograft in avian embryos"

**Supplemental Data**

**Supplemental Tables** .......... 2

**Supplemental Figures** .......... 7

### Supplemental Tables

Supplemental Table 1: Characteristics of the patients included in the study.

| Patient Number | Anatomical localization of the cryopreserved lymph node | Age at diagnosis | Sex | Histological grade | FLIPI score | Time elapsed between treatment initiation and event or last follow-up (months) | Event (relapse or death) | Analyses realized in this study | | |
|---|---|---|---|---|---|---|---|---|---|---|
| | | | | | | | | scRNA-seq | RCHOP imaging | RCHOP+Venetoclax imaging |
| 1 | axillary | 58 | M | 2 | 1 | 13 | Yes | X | X | |
| 2 | parotid | 53 | F | 2 | 0 | 22 | Yes | X | X | |
| 3 | cervical | 56 | M | 1 | 1 | 7 | Yes | X | X | |
| 4 | axillary | 47 | M | 2 | 1 | 185 | No | X | X | |
| 5 | sus-clavicular | 53 | M | 2 | 1 | 164 | No | X | X | |
| 6 | inguinal | 56 | M | 3a | 0 | 138 | No | | X | |
| 7 | inguinal | 44 | F | 1-2 | 0 | 185 | No | X | X | |
| 8 | sub-maxillary | 75 | F | 1-2 | 4 | 8 | Yes | | X | |
| 9 | inguinal | 60 | F | 2 | 3 | 9 | Yes | | X | |
| 10 | mesenteric | 60 | M | 2 | 1 | 7 | Yes | X | X | |
| 11 | axillary | 51 | M | 3a | 2 | 168 | No | X | X | |
| 12 | inguinal | 47 | M | 2 | 2 | 85 | Yes | X | X | |
| 13 | axillary | 79 | M | 1-2 | 4 | 4 | Yes | | X | |
| 14 | inguinal | 52 | M | 1-2 | 2 | 98 | No | | X | |
| 15 | inguinal | 61 | M | 1 | 3 | 31 | No | X | X | |
| 16 | cervical | 59 | F | 1 | 2 | 157 | Yes | X | X | |
| 17 | inguinal | 91 | M | 1-2 | 4 | 34 | Yes | | X | X |
| 18 | inguinal | 52 | M | 1-2 | 2 | 12 | Yes | X | X | X |
| 19 | axillary | 41 | F | 1-2 | 1 | 45 | No | | X | X |
| 20 | axillary | 61 | M | 1 | 2 | 39 | No | X | X | X |
| 21 | inguinal | 46 | M | 1-2 | 3 | 42 | No | | | X |
| 22 | cervical | 82 | M | 1-2 | 4 | 24 | No | | | X |
| 23 | inguinal | 49 | F | 2 | 2 | 125 | No | X | | |

Supplemental Table 2: Raw data of FL-AVI-PDX for percentage of tumor intake and volume reduction after RCHOP exposure.

| Patient Number | Responder status | % Chick embryos viability (24H post-implantation) | % tumor intake | Number of embryos per condition | | Tumor volume reduction (%) Excipient VS RCHOP |
|---|---|---|---|---|---|---|
| | | | | Excipient | RCHOP | |
| 1 | Poor responder | 88.52% | 100% | 13 | 11 | 11% |
| 2 | Poor responder | 91.25% | 92.50% | 18 | 19 | 9% |
| 3 | Poor responder | 82.20% | 100% | 8 | 7 | 18% |
| 4 | Good responder | 85.78% | 88% | 8 | 8 | 75% |
| 5 | Good responder | 77.32% | 100% | 18 | 12 | 25% |
| 6 | Good responder | 75.70% | 86% | 11 | 12 | 35% |
| 7 | Good responder | 81.90% | 91% | 11 | 16 | 18% |
| 8 | Poor responder | 96.37% | 100% | 11 | 12 | +17% |
| 9 | Poor responder | 92.66% | 100% | 11 | 13 | +15% |
| 10 | Poor responder | 93.45% | 100% | 12 | 10 | +13% |
| 11 | Good responder | 94.28% | 97.70% | 18 | 22 | 42% |
| 12 | Good responder | 88.09% | 100% | 7 | 9 | 33% |
| 13 | Poor responder | 95.74% | 100% | 13 | 13 | +31% |
| 14 | Good responder | 76.25% | 100% | 12 | 18 | 28% |
| 15 | Good responder | 89.90% | 100% | 15 | 14 | 20% |
| 16 | Good responder | 93.37% | 100% | 13 | 12 | 55% |
| 17 | Poor responder | 92.94% | 100% | 17 | 9 | 13% |
| 18 | Poor responder | 88.20% | 97.70% | 12 | 12 | 8.5% |
| 19 | Good responder | 92.70% | 100% | 13 | 10 | 26% |
| 20 | Good responder | 91.06% | 100% | 14 | 10 | 26% |

Supplemental Table 3: Number of cells for each patient and condition analyzed by scRNAseq in this study.

| Patient Number | Number of total cells | Number of cells (absolute number) | | | | | | Number of reads per cell |
|---|---|---|---|---|---|---|---|---|
| | | Pre-graft condition | | Vehicle condition | | RCHOP condition | | |
| | | Passed QC | Active cycling cells | Passed QC | Active cycling cells | Passed QC | Active cycling cells | |
| 1 | 2927 | 1540 | 971 | 696 | 391 | 691 | 383 | 128.371 |
| 2 | 7276 | 3133 | 1943 | 2089 | 1249 | 2054 | 1246 | 66.382 |
| 23 | 2133 | 1360 | 821 | 418 | 233 | 355 | 224 | 54.441 |
| 3 | 893 | 679 | 399 | 128 | 83 | 86 | 50 | 86.870 |
| 7 | 3434 | 1165 | 648 | 1111 | 580 | 1158 | 619 | 81.170 |
| 4 | 1287 | 1103 | 598 | 124 | 50 | 60 | 37 | 97.729 |
| 5 | 8439 | 3045 | 1996 | 2776 | 1680 | 2618 | 1618 | 43.852 |
| 10 | 1166 | 657 | 474 | 305 | 213 | 204 | 142 | 97.715 |
| 11 | 2726 | 1610 | 1138 | 639 | 449 | 477 | 313 | 65.648 |
| 15 | 1759 | 817 | 547 | 513 | 354 | 429 | 283 | 84.428 |
| 12 | 941 | 354 | 278 | 279 | 191 | 308 | 206 | 256.765 |
| 16 | 736 | 401 | 289 | 240 | 136 | 95 | 46 | 146.841 |
| 18 | 9344 | 3532 | 2645 | 2565 | 1718 | 3247 | 2120 | 25.125 |
| 20 | 2192 | 980 | 513 | 639 | 343 | 573 | 325 | 53.846 |

Supplemental Table 4: Doses of individual drugs of the RCHOP combination for *in ovo* experimentation and raw data of MTD experimental determination.

| | RCHOP-combination | | | | | Viability | |
|---|---|---|---|---|---|---|---|
| Dose-ranging | Rituximab (mg/kg) | Cyclophosphamide (endoxan) (mg/kg) | Hydroxyadriamycine (mg/kg) | Vincristine (mg/kg) | Prednisone (mg/kg) | Percentage of viability (48h) | Number of embryos analyzed |
| Excipient (Equal volume) | NaCl 0.9% + DMSO | NaCl 0.9% + DMSO | NaCl 0.9% + DMSO | NaCl 0.9% + DMSO | NaCl 0.9% + DMSO | 93.3 | 15 |
| C1 | 5.13 | 10.25 | 0.69 | 0.02 | 0.54 | 86.6 | 15 |
| C1*2 | 10.25 | 20.5 | 1.38 | 0.04 | 1.075 | 60 | 15 |
| C1*4 | 20.5 | 41 | 2.76 | 0.08 | 2.15 | 26.6 | 15 |
| C1*8 | 41 | 82 | 5.52 | 0.16 | 4.3 | 13.3 | 15 |

Supplemental Table 5: RCHOP signature genes (n= 21) with associated logFC and FDR in single cell and pseudo-bulk analysis.

| Genes | logFC | | FDR | |
|---|---|---|---|---|
| | Single cell | Pseudo-bulk | Single cell | Pseudo-bulk |
| RPS19 | 0.275 | 0.269 | 2.50e-183 | 4.40e-02 |
| BAX | 0.467 | 0.605 | 3.60e-165 | 2.13e-7 |
| PSMB4 | 0.387 | 0.535 | 1.25e-145 | 7.09e-8 |
| PHPT1 | 0.453 | 0.698 | 2.85e-135 | 1.36e-05 |
| RPS27L | 0.377 | 0.478 | 5.50e-127 | 4.43e-05 |
| SRSF3 | 0.263 | 0.325 | 1.57e-61 | 1.89e-02 |
| HIST1H2BK | 0.412 | 0.794 | 1.17e-58 | 6.29e-9 |
| TRIM22 | 0.269 | 0.478 | 1.24e-56 | 1.31e-06 |
| FDXR | 0.227 | 0.956 | 3.78e-55 | 1.18e-06 |
| MRFAP1 | 0.25 | 0.476 | 4.62e-55 | 7.04e-05 |
| LY86 | 0.226 | 0.347 | 1.92e-46 | 3.57e-04 |
| PVT1 | 0.203 | 0.578 | 6.00e-44 | 2.83e-04 |
| DDB2 | 0.175 | 0.509 | 1.13e-31 | 1.71e-04 |
| AEN | 0.165 | 0.635 | 6.07e-29 | 6.08e-05 |
| PQBP1 | 0.16 | 0.28 | 1.87e-22 | 2.92e-02 |
| CD70 | 0.217 | 0.527 | 7.77e-22 | 3.99e-04 |
| BBC3 | 0.173 | 0.613 | 1.63e-20 | 7.04e-05 |
| HNRNPH1 | 0.159 | 0.308 | 3.25e-19 | 1.14e-02 |
| SRSF2 | 0.163 | 0.294 | 8.33e-19 | 1.67e-02 |
| CDKN1A | 0.193 | 0.401 | 2.01e-13 | 2.67e-02 |
| MT2A | 0.154 | 0.595 | 3.91e-3 | 1.86e-02 |

Supplemental Table 6: Raw data of venetoclax MTD experimental determination when used alone or in combination with RCHOP MTD.

Toxicity assay Venetoclax alone

| | Standard criteria | | Number of embryo analyzed | |
|---|---|---|---|---|
| Treatment | Percentage of viability (48h) | Mean BSA. (cm² +/- SEM) | Viability | BSA |
| Excipient (Equal volume) NaCl 0.9% + Tween 80 | 80.75% | 0.935 ± 0.025 | 26 | 21 |
| Venetoclax 2.12 mg/kg | 76.92% | 0.906 ± 0.04 | 13 | 10 |
| Venetoclax 6.12 mg/kg | 76.92% | 0.939 ± 0.028 | 13 | 18 |
| Venetoclax 19.14 mg/kg | 84.61% | 0.952 ± 0.026 | 13 | 11 |
| Venetoclax 57.43 mg/kg | 69.23% | 0.8538 ± 0.036 | 13 | 9 |
| Venetoclax 172.29 mg/kg | 87.82% | 0.8615 ± 0.041 | 26 | 17 |
| Venetoclax 516.93 mg/kg | 46.15% | 0.9035 ± 0.051 | 13 | 6 |
| Venetoclax 1550 mg/kg | 46.15% | 0.784 ± 0.035 | 13 | 6 |

Toxicity assay Venetoclax in combination with R-CHOP

| | | Standard criteria | | Number of embryo analyzed | |
|---|---|---|---|---|---|
| Treatment | | Percentage of viability (48h) | Mean BSA (cm² +/- SEM) | Viability | BSA |
| + RCHOP Dose Ranging 1 (MTD) | Excipient (Equal volume) NaCl 0.9% + Tween 80 | 100% | 0.8072 ± 0.04 | 13 | 12 |
| | Venetoclax 6.12 mg/kg | 76.92% | 1.107 ± 0.033 | 13 | 10 |
| | Venetoclax 19.14 mg/kg | 92.30% | 1.032 ± 0.038 | 13 | 12 |
| | Venetoclax 57.43 mg/kg | 69.23% | 0.982 ± 0.028 | 13 | 9 |
| | Venetoclax 172.29 mg/kg | 46.15% | | 13 | |

### Supplemental Figures

**Supp. Figure 1: Establishment of the patient-derived xenograft avian model of follicular lymphoma (FL-AVI-PDX).**

**(A)** Flow cytometry gating strategy for human CD45+ CFSE+ cell sorting. The maximum number of cells were sorted for the post-graft condition, and an equivalent number of cells were sorted for the pre-graft fraction to avoid imbalance in the number of cells per condition. **(B)** Expression of the dissociation signature by cells after graft with three different protocols. **(C)** Percentage of cells expressing IGL or IGK light chain in each sample (n= 13). **(D)** Principal component analysis (PCA) representation of the pre- and post-graft scRNAseq data. Individual cells are labeled according to their phenotype (left panel) or to experimental condition (right panel). **(E)** Schematic representation of the analytical strategy used to identify the differentially expressed gene list based on single cell and pseudo-bulk methods.

**Supp. Figure 2: FL-AVI-PDX captures clinical heterogeneity of response to RCHOP.**

**(A)** Survival rate and mean body surface area (BSA) of avian embryos injected with increasing doses of RCHOP combination. The maximum tolerated dose (MTD) is the higher dose with no impact on BSA and a survival rate greater than 80%. Data are expressed as mean +/- SEM, * $p<0.05$ (Mann-Whitney test). **(B)** Comparison of the pharmacokinetics of cyclophosphamide, hydroxyadriamycin and vincristine in human (data from [14]) (dotted line) and in avian embryos (continuous line). **(C)** Kaplan-Meier progression free survival curve of the whole cohort (n= 20).

**Supp. Figure 3: FL-AVI-PDX reveals a transcriptomic signature of response to RCHOP in primary FL cells**

**(A-B)** Quantitative RT-PCR assessment of the relative expression of BAX, RPS27L, SRSF3 and RPS19 (included in the RCHOP signature) normalized on GAPDH expression in the SUDHL-4 cell line exposed to vehicle or RCHOP overnight **(A)** or to each of the five compounds individually **(B)**. **(C)** Sample of origin of cells of cluster 8 (n= 590 single cells).

**Supp. Figure 4**: **BAX is a targetable effector of RCHOP in FL**

**(A)** Western blot analysis of BAX protein levels in SUDHL-4 cells ($BAX^{-/-}$ with two different sgRNA targeting BAX or $BAX^{WT}$ with a non-target sgRNA). **(B)** Schematic representation of the mechanism of action of venetoclax. **(C)** Proportion of viable (annexin V negative) SUDHL-4 cells ($BAX^{-/-}$ with two different sgRNA targeting BAX or $BAX^{WT}$ with a non-target sgRNA) after 48h exposure to increasing doses of venetoclax (µM). Error bars indicate SD. *p Val<0.05 (Wilcoxon test). **(D)** Survival rate and mean body surface area (BSA) of avian embryos injected with increasing doses of venetoclax. The maximum tolerated dose (MTD) is the higher dose with no impact on BSA and a survival rate greater than 80%. Data are expressed as mean +/- SEM, **p Val <0.005, ns not significant (Mann-Whitney test). **(E)** Survival rate and mean body surface area (BSA) of avian embryos injected with increasing doses of venetoclax combined with the MTD of RCHOP. **p Val <0.005, ***p Val <0.0001 (Mann-Whitney test).

Supp. Figure 1

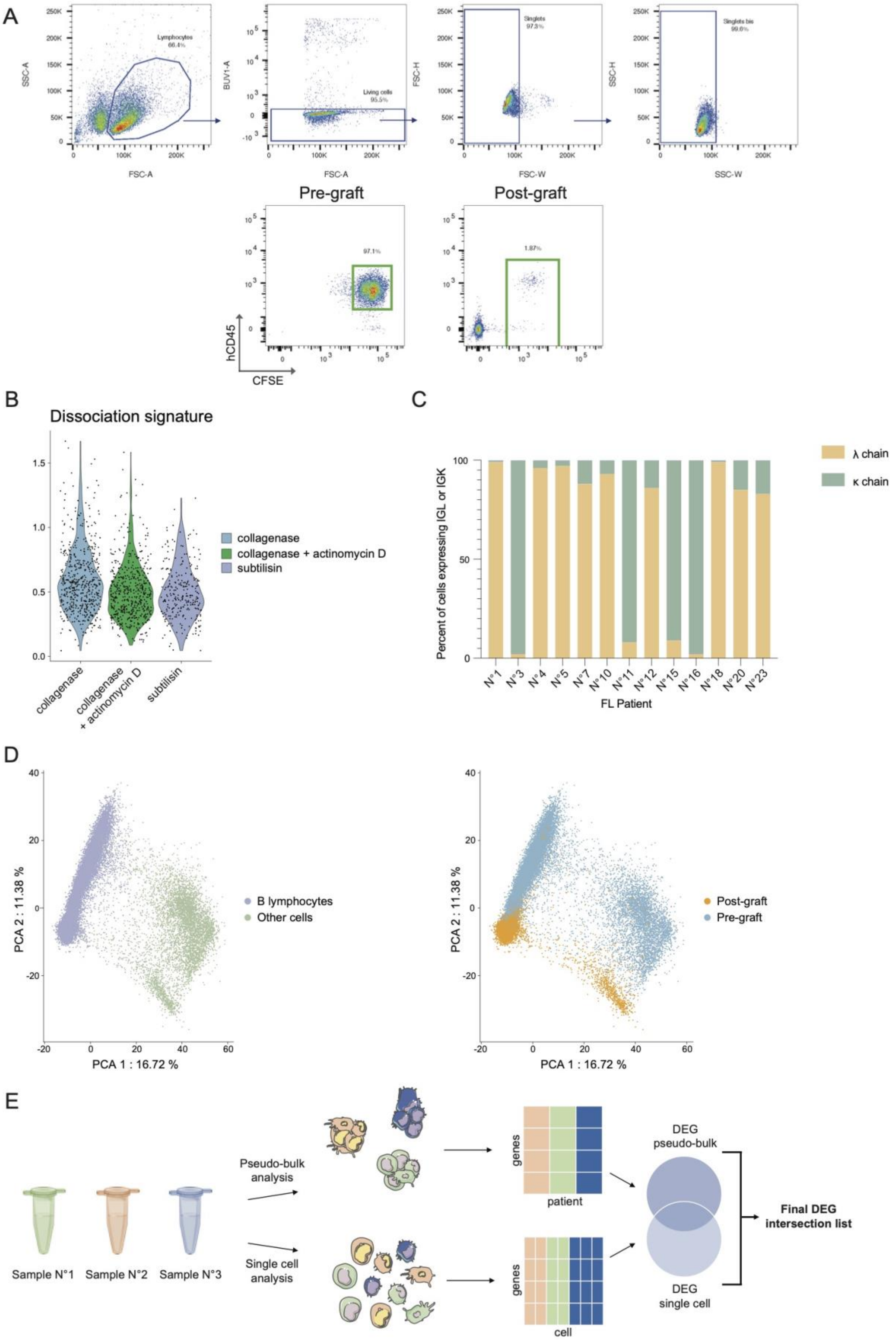

Supp. Figure 2

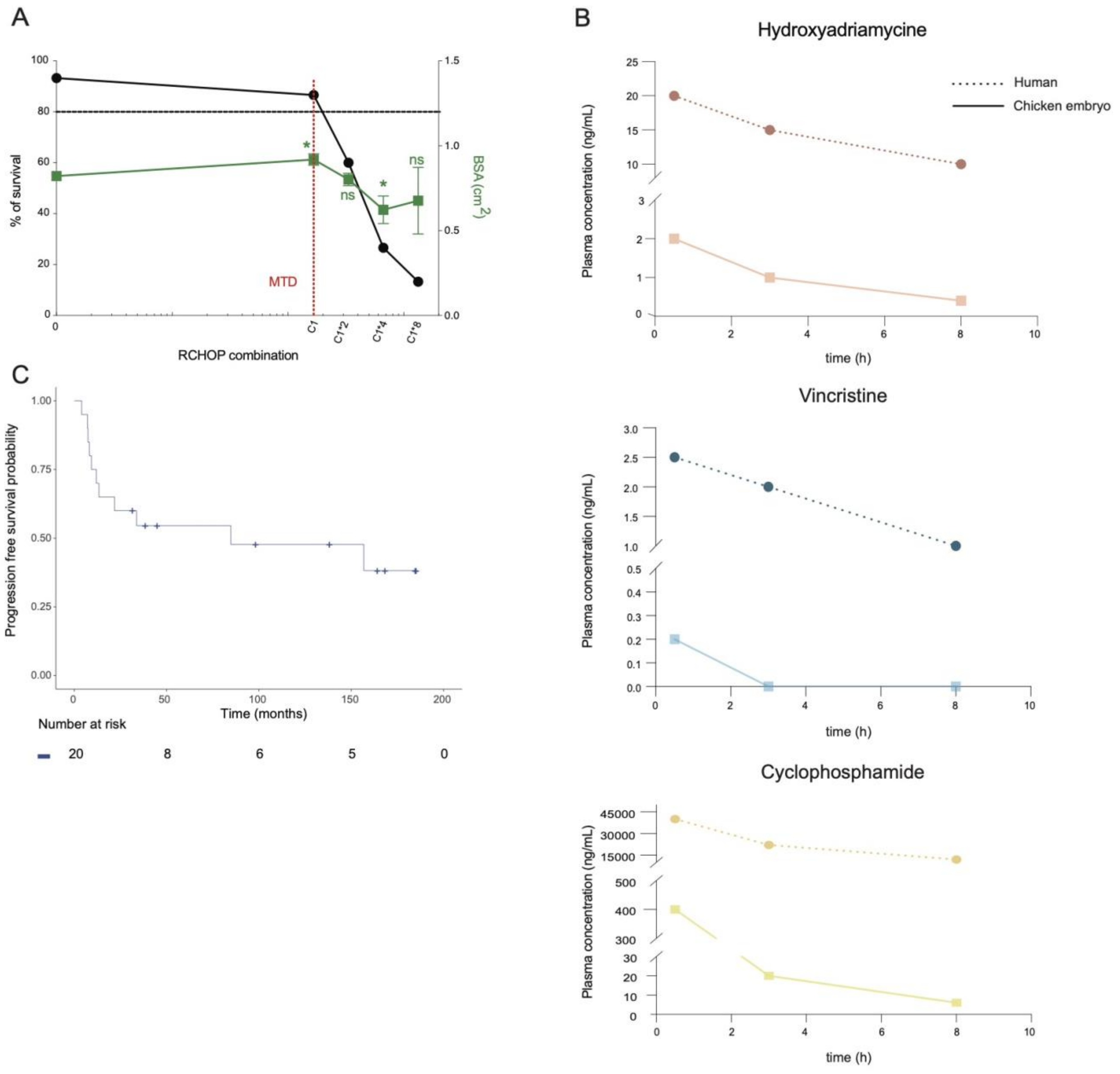

Supp. Figure 3

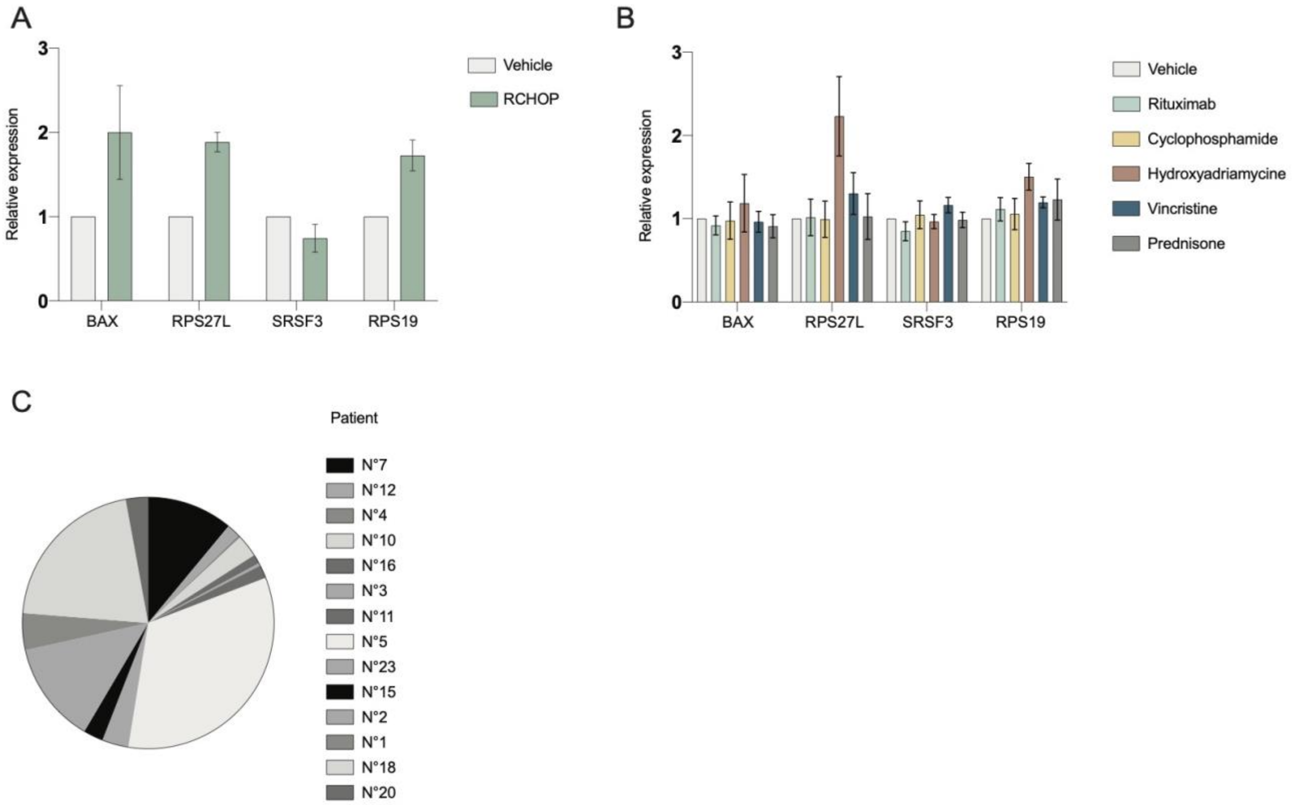

Supp. Figure 4

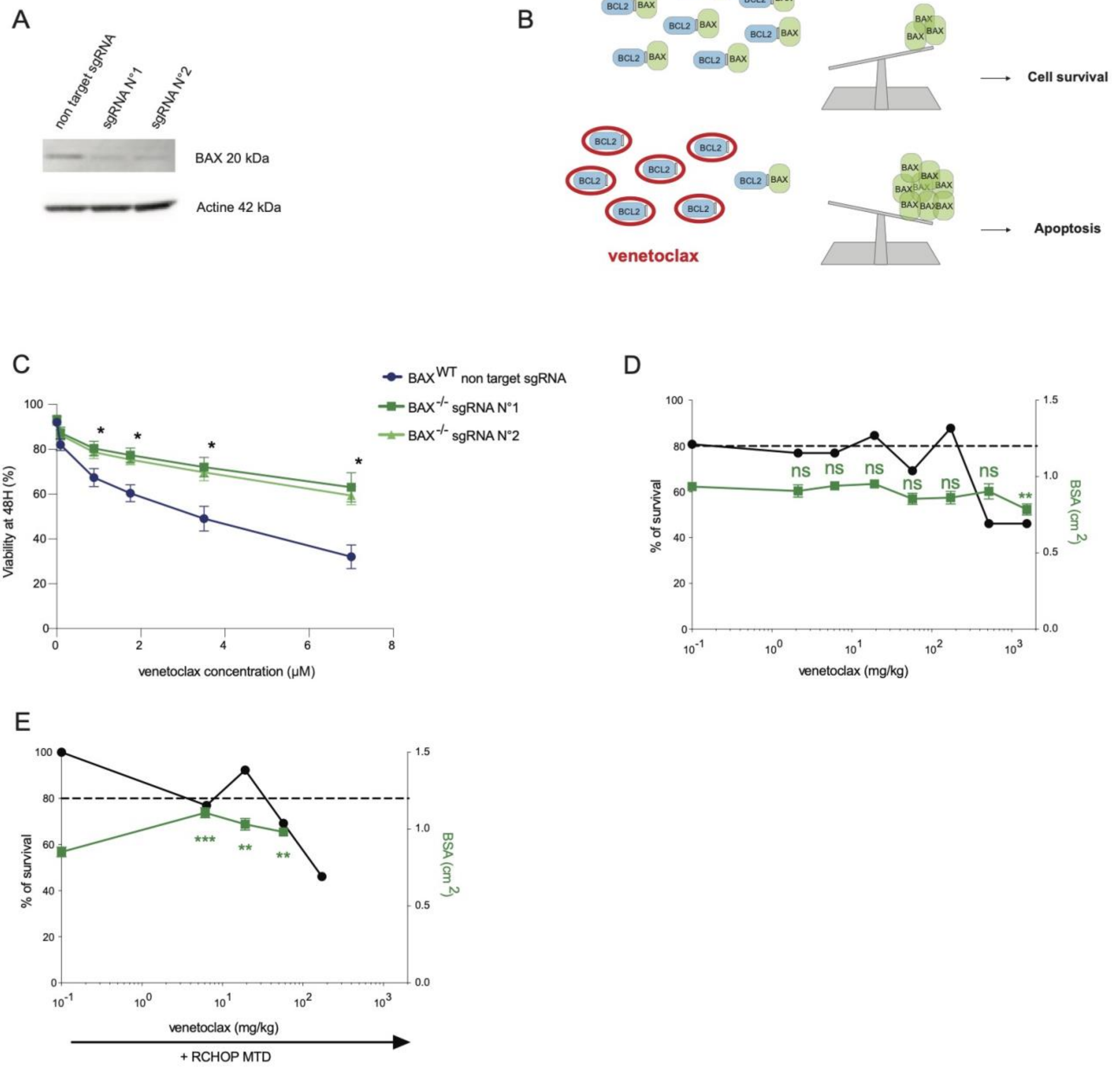
